# Impact of Interval Censoring on Data Accuracy and Machine Learning Performance in Biological High-Throughput Screening

**DOI:** 10.1101/2024.09.25.615059

**Authors:** Vanni Doffini, Michael A. Nash

**Affiliations:** University of Basel; ETH Zurich; Swiss Nanoscience Institute

## Abstract

High-throughput screening (HTS) combined with deep mutational scanning (DMS) and next-generation DNA sequencing (NGS) have great potential to accelerate discovery and optimization of biological therapeutics. Typical workflows involve generation of a mutagenized variant library, screening/selection of variants based on phenotypic fitness, and comprehensive analysis of binned variant populations by NGS. However, in such cases, the HTS data are subject to interval censoring, where each fitness value is calculated based on the assignment of variants to bins. Such censoring leads to increased uncertainty, which can impact data accuracy and, consequently, the performance of machine learning (ML) algorithms tasked with predicting sequence-fitness pairings. Here, we investigated the impact of interval censoring on data quality and ML performance in biological HTS experiments. We theoretically analyzed the impact of data censoring and propose a dimensionless number, the *Ratio of Discretization* (*R*_*D*_), to assist in optimizing HTS parameters such as the bin width and the sampling size. This approach can be used to minimize errors in fitness prediction by ML and to improve the reliability of these methods. These findings are not limited to biological HTS techniques and can be applied to other systems where interval censoring is an advantageous measurement strategy.

## 1 Introduction

High-throughput screening (HTS) [43, 1, 27, 55, 6] has profoundly transformed the molecular engineering field by enabling the rapid assessment of thousands to millions of protein variants for key properties such as enzymatic activity and ligand binding. When coupled with advanced methodologies such as deep mutational scanning (DMS) [18, 42, 24] and next-generation DNA sequencing (NGS) [17, 40, 19, 16], HTS becomes a powerful tool for the identification and evolution of novel therapeutics from vast mutagenized libraries. This integrated approach not only accelerates the discovery process but also reduces associated costs, making it indispensable for modern biotechnology.

A typical workflow for variant discovery is based on fluorescent-activated single-cell sorting (FACS) [13, 35, 46] coupled with NGS, and includes three main steps [47, 48]: (i) generation of a mutagenized library of protein variants containing sequence alterations; (ii) single-cell sorting of the variant pool into different bins based on their fluorescence intensity by FACS, and (iii) analysis of the populations in each bin using NGS. In this case, the screening assay is set up such that the fluorescence intensity of the cell reports on the relevant phenotype, for example, by binding to a fluorescently labeled ligand [22]. This process establishes a genotype-phenotype relationship for each mutant screened and enables the library to be sorted according to the desired fitness trait. Subsequently, the best variants can be propagated to subsequent screening rounds and the procedure repeated to further optimize the selected sequences in a process known as directed evolution (DE) [36, 4, 51]. This approach has proven successful in improving a range of phenotypic properties, such as enzymatic activity [9] and efficiency [39], ligand binding affinity [7, 25], and thermal stability [5].

Modern variations on this technique [38, 53, 33] have sought to leverage the enormous datasets generated by DMS in order to train machine learning (ML) models to navigate and predict the fitness landscape *in silico* [10, 45, 37]. If such an approach were successful, reliable and broadly generalizable, it would enable efficient multi-objective variant optimization [14] and save significant time and cost, thereby enhancing the efficiency and scope of protein engineering for human therapies. Indeed, ML has been employed to predict beneficial mutations, reduce the experimental search space, and design proteins with desired properties more efficiently [52, 30]. For example, ML models have been applied to enzyme engineering [41] and to predict and design antibodies with high affinity for their targets [28, 44]. Furthermore, ML is not limited to mutagenized data but can also be utilized to generate de-novo proteins [50, 26, 54, 3]. However, the field is still in its infancy. Many challenges exist regarding optimal encoding strategies, algorithm classes, scalability and efficiency, and the literature is plagued by a survivor bias of the successful methods implemented within a limited context.

One challenge that has thus far received only limited attention is that datasets obtained via FACS/NGS assays typically classify variants into specific bins based on fitness. The precise value of each point, therefore, remains unknown and resides only within the defined bin boundaries. This phenomenon, known in statistics as interval censoring, introduces significant uncertainty into the dataset and makes the outcome highly dependent on the gating strategy used to define sorting bins. Further complicating the picture is the fact that each variant is represented by multiple cells and therefore generates multiple data points. Due to intrinsic biological variability and experimental noise, these cellular replicates of the variants frequently fall into different bins, resulting in a censored distribution of fitness values. A prior study focused on inferring protein fitness properties from the classification of variants in two bins [8], but did not address larger numbers of bins or how the number of collected samples influences measured fitness.

Additionally, from a purely practical perspective, it is important to consider not only censoring error but also its associated costs. Enhancing data resolution by increasing the number of gates (i.e., decrease the gate width) and collecting more measurements to achieve a more accurate fitness distribution and improve reproducibility inevitably increases experimental costs. In most cases, the number of gates will be limited by the choice of hardware (i.e., FACS instrument) and is generally limited to low numbers in a single sorting run (e.g., typically 4-channel sorting). Therefore, achieving an optimal trade-off between the various factors is crucial for efficient and cost-effective implementation of HTS in research.

The ultimate goal is to ensure that realistically-sampled censored data are truly representative of each variant-specific theoretically continuous distribution of fitness values. This can be achieved by minimizing the divergence between a specific measure derived from the censored data (i.e., the censored averages 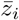) and the ideal data (i.e., the continuous means *µ*_*y,i*_) for each variant in the library. Moreover, ML presents an additional layer of complexity. In fact, ML models are trained on censored data 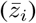, but their predictions should ideally represent the theoretical fitness values (*µ*_*y,i*_).

A critical aspect of ML in this setting is its ability to learn efficiently from training examples, which can be quantified using learning curves (LCs) [11, 29], a popular concept in ML applied to quantum physical chemistry [20]. LCs reflect how a model’s predictive performance improves as more data points are added to the training set, and typically follow a monotonically decreasing trend [49]. Historically, LCs have been observed to follow a power-law decay [2], usually depicted on a log-log scale. However, recent work on ML of protein sequences [15] suggests that when the training set is scaled to include variants with increasing numbers of mutations and normalized using a combinatorial scale, the resulting LCs show discontinuities and different decay behaviors. These are referred to as “mutant-based normalized” LCs.

Although interval censoring has been studied extensively in mathematics [56], with solutions such as Sheppard’s correction for statistical moments [23, 12], these solutions often assume a theoretical scenario involving a single probability distribution with infinite samples discretized by narrow gates. The HTS scenario, however, involves the censoring of multiple distributions, finite sampling, and broader gating strategies, which can result in all samples from a variant being enclosed within a single bin. Furthermore, the impact of interval-censored HTS data on ML performance remains largely unexplored.

To address this knowledge gap, we present a comprehensive study on the effects of interval censoring in HTS experiments on data accuracy and ML performance. After introducing the HTS parameters and the parallels between our approach and a hypothetical experimental scenario, we present a theoretical exploration of interval censoring, followed by an in-depth analysis of simulated HTS data. We then assess the performance of ML models trained on censored data using mutant-based normalized LCs, highlighting challenges, and proposing solutions for reconstructing continuous fitness distributions from censored data. Our findings indicate that optimizing HTS parameters, such as gate widths and sample sizes, can significantly reduce errors and enhance the learning efficiency of ML models. We propose the introduction of a dimensionless number, *R*_*D*_, to guide these optimizations, ensuring a balance between minimizing censoring errors and maintaining experimental feasibility. This study aims to offer practical guidelines for researchers, ultimately improving the accuracy and efficiency of HTS methodologies.

## 2 Results

### 2.1 Parameter Set and System Overview

HTS experiments based on FACS/NGS are influenced by several parameters (Fig. 1A, Tab. S2). In this study, we focused on the effect of (i) gate width (*h*); (ii) gate positioning (*z*_0_ or *h*_0_); (iii) the average number of samples acquired 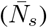; and (iv) the distribution of mutagenized variants in the screened library (*p*_*WT*_). Additionally, we kept the following parameters constant: the wild-type (*WT*) sequence and its length (*n*), the mutation vocabulary (*v*), the functional form of the continuous fitness landscape (*f* (*·*)), along with the corresponding variability in fitness measurements (*σ*_*y*_). Even though such parameters are also crucial factors that could influence HTS outcomes, we kept them constant to simplify and focus our study.

**Figure 1.**
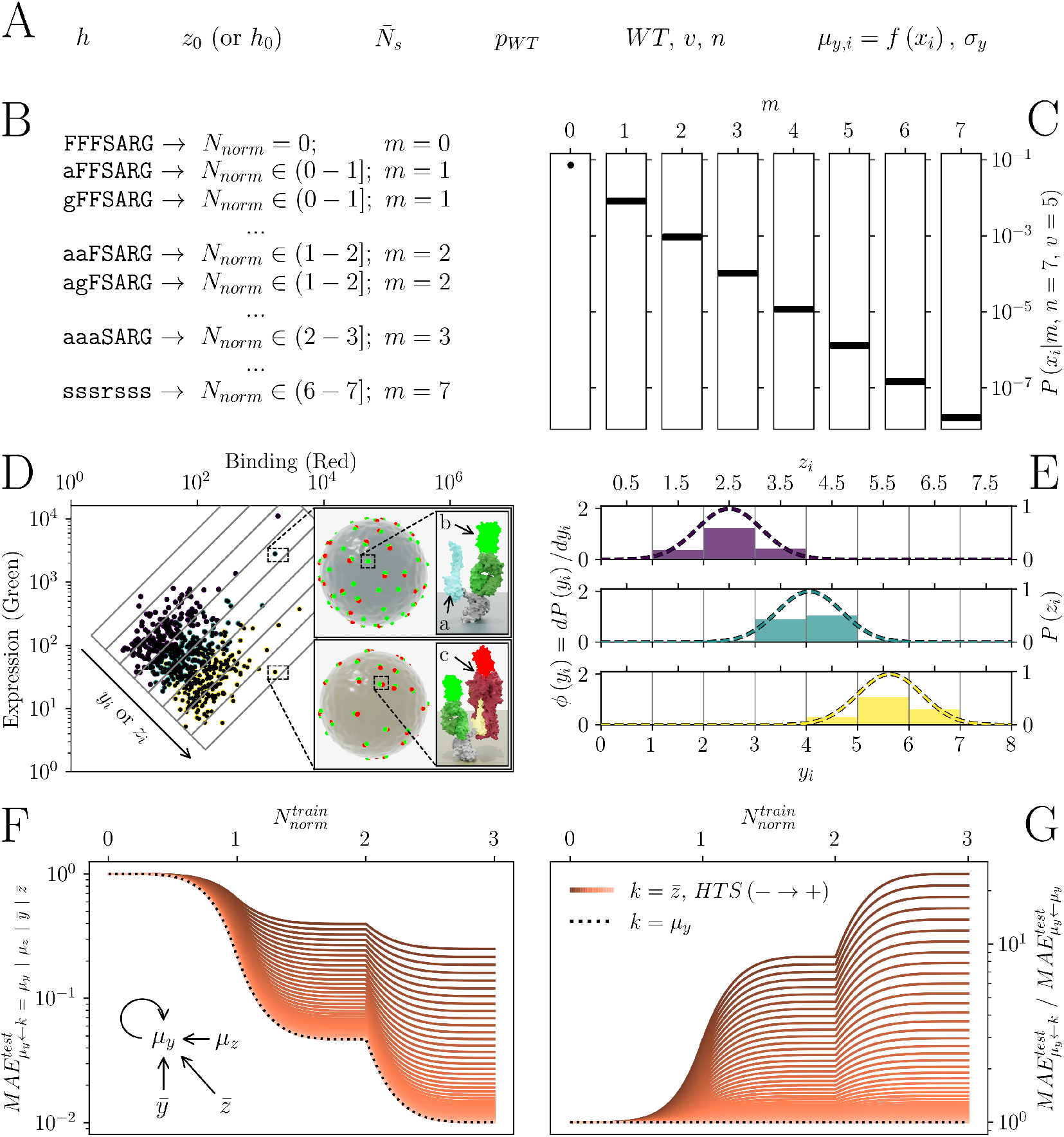
Overview of mutagenized library generation, FACS-based sorting and ML evaluation with interval-censored data. (A) Key parameters influencing interval censoring in FACS experiments. (B) Design of a combinatorial variant library derived from the parent peptide sequence FFSARG, with different mutation positions leading to variant distributions. (C) Example distribution of the combinatorial library, following a binomial distribution pattern based on the mutant frequency, where the probability of retaining the wild-type sequence (*p*_*WT*_) was approximately 0.69. (D) Hypothetical FACS experiment designed to evaluate expression levels (green fluorescence) and binding affinity (red fluorescence) across variants. The first level zoom insets show two labeled cells expressing two variants (yellow and blue). The second level zoom insets show: (a) an unbound mutant peptide (blue), (b) labeled (green) expression antibody (dark green) bound to its tag (grey), and (c) labeled (red) target (dark red) bound to a mutant variant (yellow). (E) Example of population distributions of variants as determined by FACS. Dashed lines represent theoretical continuous distributions for each variant (true distributions), while the solid bars represent the binned/censored distributions (observed data). (F) Example comparison of mutant-based normalized LCs for models trained on interval-censored data 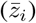 versus models trained on true continuous data (*µ*_*y,i*_), with varying HTS parameters (color gradient). The inset depicts the possible approximation/prediction-training pairings. (G) Ratio of LCs of models trained on censored averages 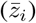 versus those trained on true means (*µ*_*y,i*_).

The type of HTS experiments we consider here typically begin with construction of a DNA library encoding the variants to be screened. This library is constructed starting from a WT sequence where mutations are introduced from a chosen vocabulary (*v*). The vocabulary refers to the set of amino acids eligible to be substituted at each position in the WT protein sequence. As a concrete example, we focus on a segment of the human fibrinogen beta chain (WT, *FFFSARG, n* = 7), and we allow mutations to be selected from the amino acids that are already present within the WT sequence (*v* = 5, amino acids: A, F, G, R and S). This peptide binds to a partner protein called SdrG from *S. epidermidis* [32]. The structure of the complex between the WT peptide and SdrG is known (PDB: 1r17), enabling us to use computationally predicted binding energy as a fitness measure for the mutagenized peptides. Binding energy was calculated via an energy force field function (EvoEF [31, 21], Supporting Information). In Fig. 1B, we present examples of mutagenized variants in the library, each shown with the corresponding number of mutations (*m*) and their associated position along the combinatorial normalized scale used in this work (*N*_*norm*_, Supporting Information).

Another critical aspect of HTS outcomes is the frequency of each variant in the library. The number of mutations (*P* (*m*)) is assumed to follow a binomial distribution, where a mutation at each position is equally likely. Under this assumption, the probability of any given variant with a known number of mutations can be calculated (*P* (*x*_*i*_ | *m*), Fig. 1C). However, this binomial distribution is just one possible model for mutation frequency. Depending on the library design strategy, other distributions may also be relevant. The binomial model is used here for simplicity, but alternative approaches could still yield similar analytical outcomes.

In an ideal experimental scenario, the DNA library is transformed into a host cell line and protein expression is induced using standard biochemical techniques. The variant library is then characterized and screened for a specific fitness of interest using FACS-based HTS. In Fig. 1D, we illustrate the hypothetical scenario for this study, where a library of mutagenized peptides is screened for binding activity against a target protein. The fitness parameter of interest is the binding affinity between the peptide variant (Fig. 1D, a) and the target protein. Each variant in the combinatorial library is displayed on the surface of a host cell (black dots), typically *E. coli* or *S. cerevisiae*, in a multivalent (i.e. many thousands of copies per cell) manner, with one variant type per cell and multiple cells per variant type. A fluorescently labeled (i.e., green) antibody (Fig. 1D, b) can bind to the displayed protein to estimate the quantity of molecular receptors (the displayed peptide) expressed on the cell surface. Subsequently, a fluorescently labeled (i.e., red) protein target (Fig. 1D, c) would be screened against the variant peptide library using FACS. Measuring both the display level and binding level is crucial to correctly normalize the binding affinity to expression levels. In such an experiment, FACS sorting gates corresponding to peptides exhibiting equal affinity (*y* or *z*) with the target protein can be drawn obliquely (Fig. 1D). The cells contained within one of these slanted bins are then sorted by the FACS instrument, and processed by NGS to obtain the bin-wise interval-censored distribution of each variant (Fig. 1E).

An efficient data analysis algorithm would reconstruct the continuous distribution of each variant from its censored distribution and ultimately obtain an accurate estimate of true fitness values (i.e., the true means of the continuous distributions, *µ*_*y,i*_) from the binned data (the averages of the censored distributions, 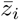). Since we generated our datasets via simulations rather than experiments, we had access to two additional approximations, namely the censored means (*µ*_*z,i*_) and the continuous averages 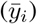. Later in this study, we used these values to evaluate the divergence of each approximation level (Tab. S3) from *µ*_*y,i*_ by calculating their mean absolute error (MAEs) using different binning and sorting strategies. Subsequently, we utilized LCs to evaluate ML performance on predicting *µ*_*y,i*_ from models trained with data coming from the different approximations (*µ*_*y,i*_, *µ*_*z,i*_, 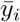 and 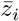) for different HTS parameter sets. Such LCs were normalized using a combinatorial scaling based on the number of mutations (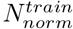, Supporting Information). Moreover, we also report the ratio of LCs (Fig. 1G) by normalizing LCs to the ones corresponding to the same models trained on continuous means (*µ*_*y,i*_).

### 2.2 Theoretical distribution of a single variant (*N*_*s*_ = *∞*)

We began by analyzing the impact of censoring on a single Gaussian distribution (Fig. 2A), focusing on the effects of the normalized gate width (*h/σ*_*y*_) and gate position (*z*_0_*/h*). This analysis involved measuring the difference between the means of the continuous and censored distributions (*µ*_*y*_ *−µ*_*z*_, Fig. 2B) and the censored standard deviation (*σ*_*z*_, Fig. 2C). Both quantities were normalized by the standard deviation of the continuous distribution (*σ*_*y*_).

**Figure 2.**
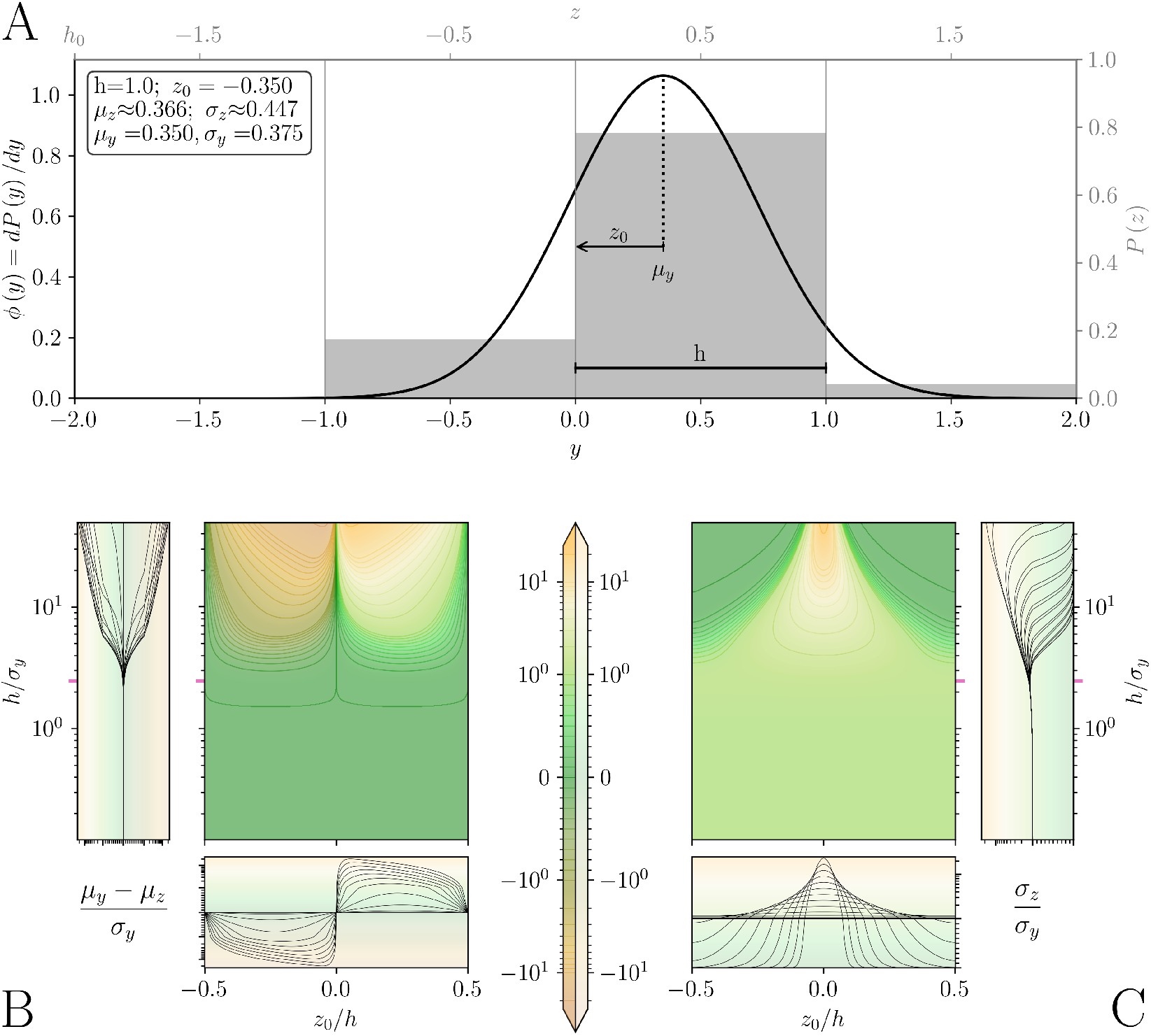
Impact of interval censoring on a single variant with ideally distributed fitness (gaussian), changing the normalized gate width (*h/σ*_*y*_) and the normalized gate position (*z*_0_*/h*). (A) Example of continuous gaussian distribution (black line) and censored distribution associated with it (grey bars). (B) Contour plots display the difference in normalized continuous mean (*µ*_*y*_*/σ*_*y*_) and censored mean (*µ*_*z*_*/σ*_*y*_). The side and bottom panels provide cross-sectional views along the x-axis (*z*_0_*/h*) and y-axis (*h/σ*_*y*_). (C) Similar to (B), but showing the ratio of censored to continuous standard deviations (*σ*_*z*_*/σ*_*y*_). The colormap is shared between panels (B) and (C).

The normalized difference in means (Fig. 2B) exhibited an antisymmetric relationship with respect to the y-axis (*z*_0_*/h* = 0). This difference converged to zero when the continuous distribution was located exactly in the middle of a gate (*z*_0_*/h* = *±* 0.5), when the population was equally divided in two parts (*z*_0_*/h* = 0), or when the gate width was below a certain threshold. The first observation was related to the fact that when the gate width was sufficiently large, the entire population could be contained within a single gate. Conversely, when the population was divided into a sufficient number of gates, the continuous and censored populations converged (Kullback–Leibler divergence approached zero, Fig. S1A), resulting in equal means.

The normalized censored standard deviation (Fig. 2C) exhibited a symmetric behavior, with a peak occurring at a normalized position of zero (*z*_0_*/h* = 0). At high gate width, the magnitude of this peak increased with the growth of *h/σ*_*y*_ following the formula for a binary discrete distribution (Eq. 1).

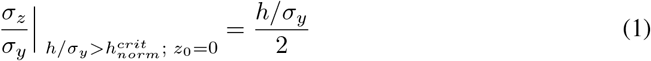

Otherwise, if the entire continuous population was contained within a single gate, the censored standard deviation collapsed to zero. This outcome did not imply that the general uncertainty was zero, but rather that the precise position within the gate could not be determined. Similarly, as observed for the difference in means, below a certain gate width threshold, the normalized censored standard deviation became independent of the gate position and converged. In this case, it did not converge to zero but to the Sheppard’s correction of the second moment (Eq. 2).

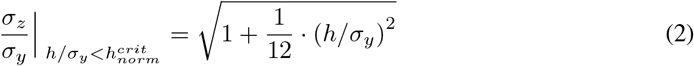

We could then define the normalized critical gate width 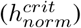 as the point at which the two previous expressions (Eq. 2 and 1) intersected (Eq. 3 and Fig. S2).

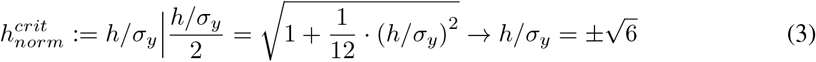

Similar observations could be made by examining the Shannon’s entropy of the censored population (Fig. S1B). In this case, 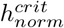 could be interpreted as the point below which the information was uniformly distributed regardless of the gate position. Beyond this point, the entropy linearly increased with the exponential decay of *h/σ*_*y*_.

### 2.3 Realistic distribution of a single variant (*N*_*s*_ *< ∞*)

We extended the previous analysis by introducing the effect of sampling. To estimate the standard error of both the continuous and censored distributions, we relied on the central limit theorem 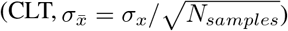. Our goal was to decouple the censoring divergence into two parts.

At first, we focused on the pure censoring error, which arises from the expected normalized absolute difference between the continuous and censored means (*µ*_*y*_ and *µ*_*z*_). We refer to this quantity as “*Bias*” or “*B*.” (Eq. 4, Fig. 3A).

**Figure 3.**
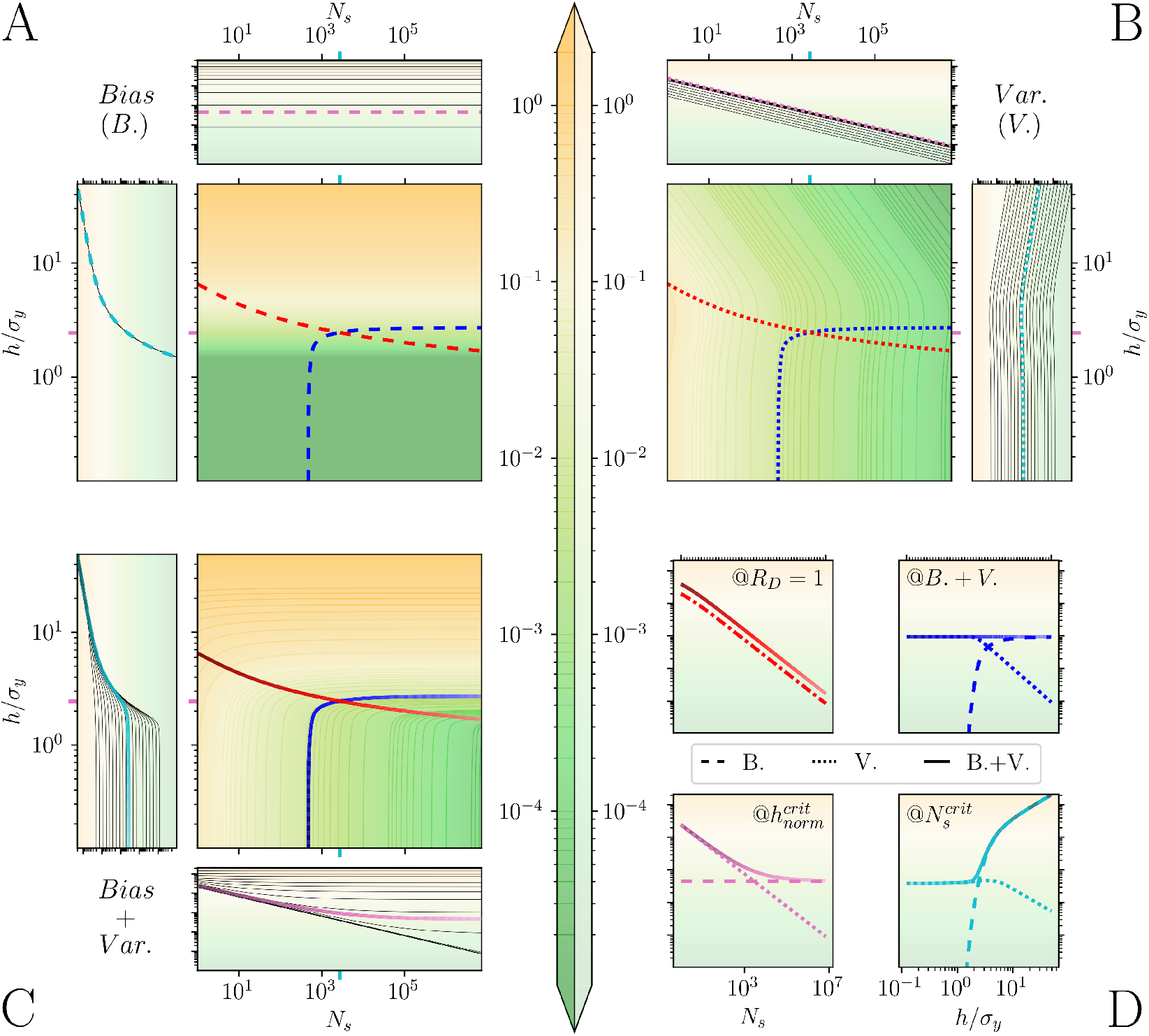
Impact of interval censoring on a single variant with realistically distributed fitness (sampled from gaussian), changing the normalized gate width (*h/σ*_*y*_) and the number of samples (*N*_*s*_). (A) Expected value of the normalized absolute difference between continuous and censored means (*Bias, B*.). (B) Expected value of the normalized absolute difference between censored mean and average (*V ar*., *V*.). (C) Sum of Bias and Variance (*Bias* + *V ar*.). (D) Contour cross-sections at different critical parameters: *R*_*D*_ = 1 (red), constant *B*. + *V*. (blue), 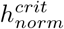 (magenta), and 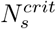 (cyan). The colormap is shared between all panels.

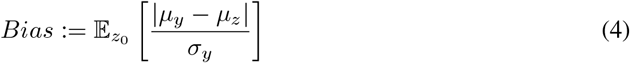

Where *z*_0_ was marginalized by calculating its expected value. *Bias* started from zero and grew exponentially with *h/σ*_*y*_, and it represented the error that one could not overcome by simply sampling the distribution more heavily (e.g., in our HTS/FACS example, by sorting more cells).

Secondly, we measured the normalized absolute difference between the censored mean and average, referring to this as “*V ar*.” or “*V*.” (*V ariance*, Eq. 5, Fig. 3B).

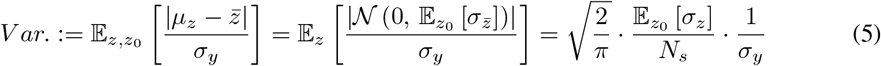

Where 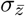 was the standard error of the censored variable accordingly to the CLT 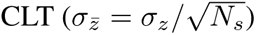 and *N*_*s*_ was the number of samples acquired.

In contrast to *Bias, V ar*. was generally dependent on both *h/σ*_*y*_ and *N*_*s*_. Nevertheless, at low gate widths 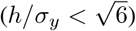, it could be approximated as dependent only on the number of samples. By our definition, *V ar*. could be seen as an estimation of the expected deviation from the biased censored mean, arising from the stochastic nature of the sampling process.

After defining *B*. and *V*., we summed them up (Fig. 3C) to estimate the maximum total divergence of the experimental censored average 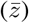 from the true continuous mean (*µ*_*y*_). Note that, due to the absolute value function, the sum of these quantities corresponded to the absolute difference between the experimental and the true values 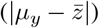 only in the worst-case scenario, where both errors had the same sign. This sum clearly showed the existence of a trade-off between *h/σ*_*y*_ and *N*_*s*_ to reach a certain maximal error, illustrated in the form of contour lines (e.g., blue line, Fig. 3C).

Finally, inspired by the chemical concept of rate-limiting reaction step, we defined a dimensionless number to measure whether the censoring error was *Bias* or *V ariance* driven (Eq. 6).

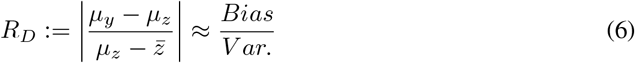

In our case, we approximated *R*_*D*_ as the ratio of *Bias* to *V ariance* by calculating its expected value with respect to *z* and *z*_0_. By setting this number to be equal to one (i.e. the point at which *B*. = *V*.) we defined a solution to the *h/σ*_*y*_ vs. *N*_*s*_ trade-off. We subsequently used this solution to define a critical number of samples by calculating its intersection with the critical gate width (Eq. 7).

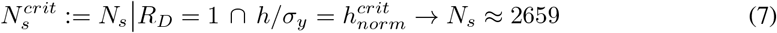

We note that, in comparison to 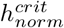, 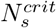 did not rely on a transition between different phases but simply on how *R*_*D*_ is calculated. Furthermore, we did not account for changes in expected experimental cost associated with decreasing the gate widths relative to scaling up the number of samples, which could be problem/experiment-specific, making the proposed solution at *R*_*D*_ = 1 potentially cost-suboptimal. Nevertheless, to show the error-optimality of *R*_*D*_ = 1, we reported *Bias, V ariance* and the sum of the two at different conditions (3D, dashed, dotted and solid lines, respectively). At *R*_*D*_ = 1 (3D, @*R*_*D*_ = 1), a power-law-like decay showed the advantage of progressing in this direction. Following a contour line (3D, @*B*. + *V*.) confirmed that *B*. + *V*. remained constant, revealing how *B*. increased non-linearly while *V*. was stable until it started to decay exponentially. Moving towards the 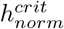 line (3D, 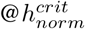) and the 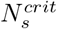 line (3D, 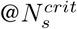) showed how *B*. + *V*. was constrained by one of the two quantities. This behavior is analogous to what could be observed in reactions when a shift in a rate-limiting regime occurs.

### 2.4 Realistic distribution of multiple variants

Until this point, our investigation focused on the distribution of fitness values of a single variant. However, our ultimate goal was to study a scenario involving a group (or pool) of distributions, specifically a pooled mutagenized library of variants displayed on cells undergoing the censoring process in a FACS experiment. Typically, the distribution of variants containing a certain number of point mutations in a combinatorial library (Fig. 1B) can be modelled using a binomial distribution. This concept could be easily extended (Eq. 8, Fig. 1C) to model the probability of a specific variant (*x*_*i*_) by knowing its number of mutations (*m*), the vocabulary size (*v*) and the length of its construct (*n*).

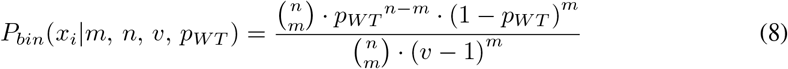

Where *p*_*WT*_ represented the probability of drawing the WT amino acid at any position, and assuming that the vocabulary included at least all amino acids present in the WT sequence.

By employing Eq. 8 along with Shannon entropy (*H*) and the Jensen Inequality, one could optimize *p*_*WT*_ to maximize the information contained in the combinatorial library (Eq. 8).

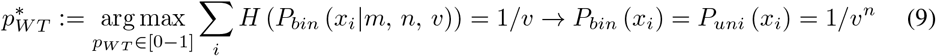

This approach corresponded to transitioning from the binomial distribution to a uniform distribution. Moreover, since it would be unfeasible to generate a library containing all possible mutations, one could use a similar approach to find a *p*_*WT*_ to maximize the information up to a specific number of mutations (*M*, Eq. 10).

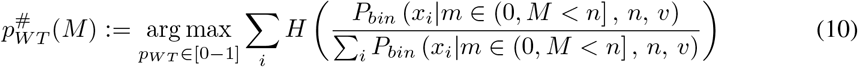

We then utilized these concepts to generate a uniformly distributed restricted library (*p*_*WT*_ = 1*/v* = 0.2, Supporting Information), containing all variant sequences with the number of mutations m ≤ 3, based on the fitness data described in our previous work [15]. We reported the mean absolute error between the continuous means (*µ*_*y,i*_ = *f*_*EvoEF*_ (*x*_*i*_)) and: (i) the censored means (*µ*_*z,i*_, Fig. S3A); (ii) the continuous averages (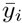, Fig. S3B); and (iii) the censored averages (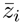, Fig. S3C). We observed the same trends previously recorded in the cases of *Bias* (Fig. 3A), *V ariance* (Fig. 3B) and *B*. + *V*. (Fig. 3C), confirming the extendibility of the concepts initially presented for a single distribution to a combinatorial library of variants.

### 2.5 ML performance on reconstructing the continuous distribution from censored training data

To evaluate the performance of ML algorithms on censored data, we utilized kernel ridge regressions (KRRs) [34] to predict the continuous means of various test sets drawn from the uniformly distributed restricted library generated in the previous section. The task for the KRR models was to reconstruct such continuous mean values of variants absent from the training set, using training data that contained varying levels of approximation. The validation and test sets comprised only continuous mean values (*µ*_*y,i*_) from variants with more than three mutations. In contrast, the training set included only variants with a maximum of three mutations and variable labels (*µ*_*y,i*_, *µ*_*z,i*_, 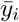, 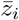).

We reported the median ratio of test MAE, calculated across different shuffles and averaged samples (Supporting Information). The ratios were calculated between models trained using censored means (Fig. S4A, *µ*_*z,i*_), continuous averages (Fig. S4B, 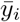), and censored averages (Fig. S4C, 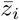), against models trained on continuous means (*µ*_*y,i*_). These results consistently demonstrated the same trends observed in the single population study (*B*., *V*., and *B*. + *V*., Fig. 3) and in the error calculation of library of variants (Fig. S3).

To further study the impact of the training dataset dimension, we plotted the mutant-based LCs at the critical values of *R*_*D*_ = 1 (Fig. 4A and S5A), at constant *B*. + *V*. (Fig. 4B and S5B), at 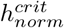 (Fig. 4C and S5C) and the 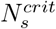 (Fig. 4D and S5D). Among such conditions, *R*_*D*_ = 1 was the only one able to achieve the same level of error as models trained with true continuous means across all possible training set sizes. This demonstrated the optimality of scaling experimental complexity according to the sets of gate widths and number of samples leading to *R*_*D*_ = 1. As expected, working at high gate widths significantly impacted the stability of learning (Fig. 4D and S5D), sometimes leading to no learning (i.e., flat LCs) or even deleterious learning, where increasing training data decreased the model performance. The data acquired from a contour line with constant *B*. + *V*. (Fig. 4B and S5B) did not demonstrate any clear advantages of high *V ar*. conditions over high *Bias* conditions. Finally, scaling up the number of samples collected per variant (Fig. 4C and S5C) consistently improved mean absolute error (MAE) at each training set size. Interestingly, at low 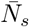, the learning curves began to lose their characteristic discontinuities, resembling a normal asymptotic power law decay.

**Figure 4.**
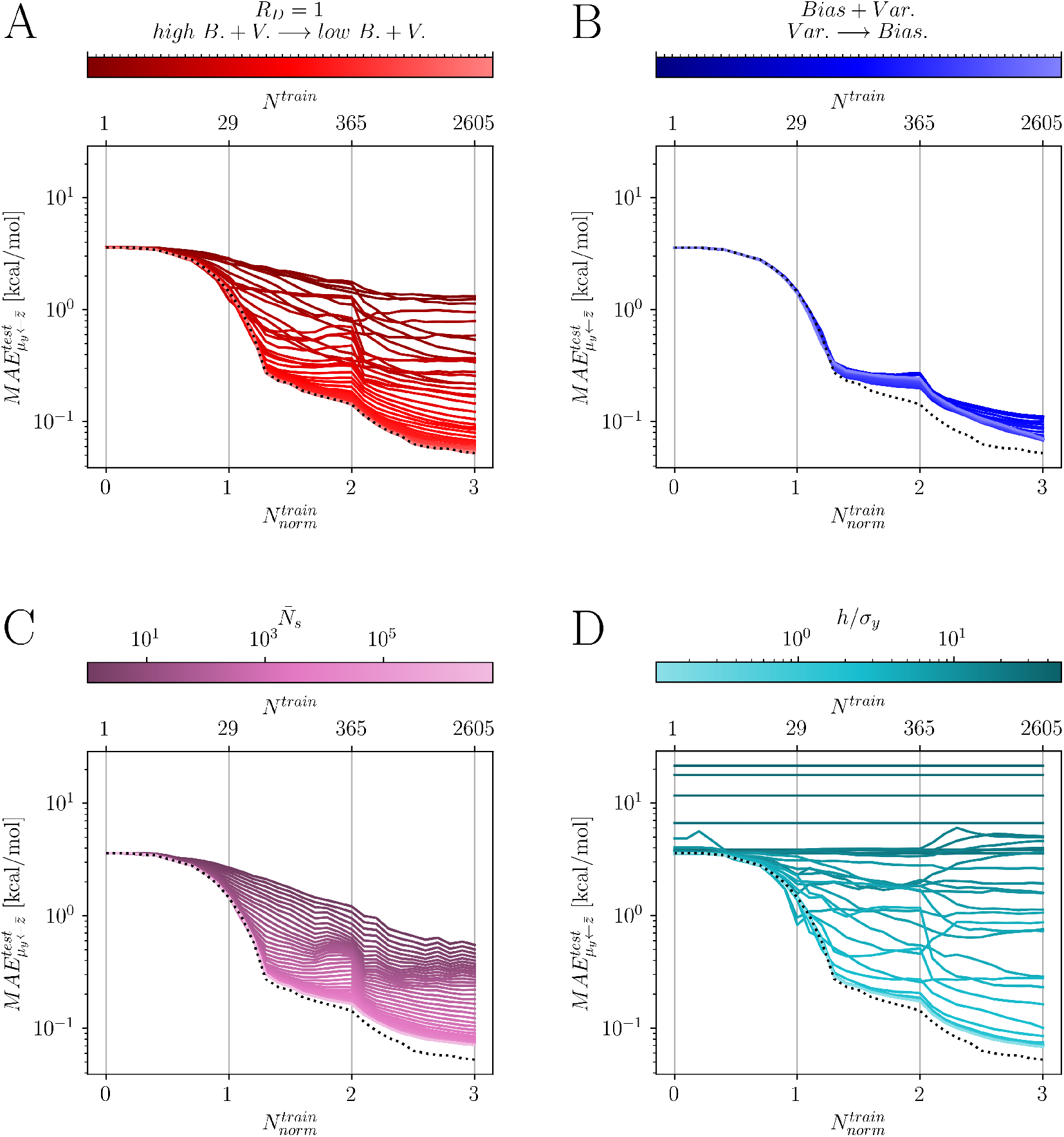
Impact of interval censoring on LCs of models trained with up to triple mutants 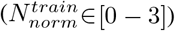 as a function of the mutant-based normalized number of training points 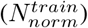, and using a uniform distributed combinatorial library (*P*_*WT*_ = 0.2). The plots illustrate the test performance under different conditions: (A) *R*_*D*_ = 1 with a gradient from high *B* + *V* to low *B* + *V* (red), (B) constant *B* + *V* with varying *B*-*V* contributions (blue), (C) constant normalized gate width 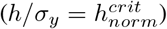 with varying average sample size (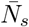, magenta), (D) constant average sample size 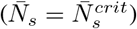 with varying normalized gate width (*h/σ*_*y*_, cyan).

In the case of a binomial distribution of variants (Fig. S6 and S7, *p*_*WT*_ *≈*0.69), similar trends were observed. However, due to the uneven distribution of samples, the uncertainty of each variant increased with the number of mutations and, consequently, with the training size. As a result, the divergence between the LCs from models trained with censored averages and those obtained with continuous means was initially lower than in the uniformly distributed case (Fig. 4 and S5). In some instances, this divergence increased with the training size until a point was reached where adding more data decreased the accuracy of the models (deleterious learning).

In general, the analysis of these LCs demonstrated that, under certain conditions, it would be more beneficial to focus on improving the acquisition parameters, such as gate width and the number of samples (to jump vertically to a different LC), rather than increasing the library size and consequently the training set size (to move right on a specific LC). This observation can significantly inform experimental design since increasing library size (and therefore screening time) can be hugely costly. The optimal gating strategy can therefore be used to maximize useful information at a given level of library complexity and screening cost.

## 3 Discussion

In this work, we investigated the impact of interval data censoring in HTS experiments where data are sorted into different bins, as is commonly the case in protein engineering campaigns relying on FACS-based single-cell sorting, next-generation DNA sequencing and machine learning. We examined increasingly complex scenarios including (i) an ideal distribution (sorting an infinite number of cells of a single variant); (ii) a sampled distribution (sorting a finite number of cells of a single variant); and multiple sampled distributions (sorting finite cells of a combinatorial variant library). In the last case, we extended our analysis by evaluating the ability of ML models to accurately reconstruct the continuous means (true fitness values) when trained on censored averages (HTS data). This approach simulated the scenario of a HTS campaign where ML would be trained on experimentally observable variants and used to predict fitness values of unseen variants not contained in the original library. As expected, it was generally possible to accurately estimate true fitness values from our synthetic HTS datasets, but at the cost of increasing the complexity of the experiments (increase number of bins / decrease bin width and increase the number of samples per variant).

We found that narrowing the sorting gate width and increasing the number of samples (i.e., the number of cells sorted) generally reduced the reconstruction error. However, modifying only one of these parameters beyond a certain threshold leads to an asymptotic behavior in the reconstruction error reduction. This asymptotic behavior was limited by either the censoring error (*Bias*) or by the sampling error (*V ariance*). We found that the preferred conditions for HTS experiments would be where neither error is dominant, and we developed a dimensionless number (*R*_*D*_) to estimate the distance of a given set of conditions from the optimal ones (*R*_*D*_ = 1). Thus, *R*_*D*_ can be utilized by experimental scientists to optimize HTS parameters, thereby minimizing the intrinsic errors arising from sampling and binning and maximizing the utility of costly experimental data.

We also quantified the critical normalized gate width 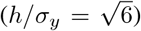 below which the differences between the censored populations (binned data) and the continuous populations began to converge to Sheppard’s correction, becoming independent of gate position. This insight can be utilized by researchers to plan their experiments, particularly in complex scenarios where *R*_*D*_ is difficult to determine or where the cost trade-off of changing the gate widths compared to the number of samples is not easily estimated. Specifically, researchers can first estimate the experimental standard deviation (*σ*_*y*_) by focusing on a single variant (e.g., WT), and then calculate the necessary gate width that should not be exceeded 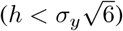.

Our analysis has certain limitations associated with it. In real-world scenarios, the number of gates required to screen an entire variant library using optimal gate widths may exceed current technological or instrument capabilities. Additionally, our assumption of a continuously normal distribution could be invalid for some systems due to various factors, such as cell non-homogeneity or skewed fitness distributions. To address the first issue, we propose to divide the experiment into multiple sorting runs with different gate offsets. For each sorting run, the number of gates should be set to the maximum allowed by the hardware, typically between 4 and 12. Gate widths should be set to ensure they cover the entire library in each screen. Subsequently, all gates should be sequentially shifted by a specific value for each sorting run. When the data from these sorting runs are combined, this approach will simulate a single sorting run with the necessary gate width. For the second issue (instrument laser/temperature fluctuations), using the median as a measure of the population fitness instead of the mean is a valid strategy. Specifically, the median should be calculated by assuming that the probability of a specific bin is continuously distributed within the gate boundaries, rather than being entirely located in the middle as in a purely discrete distribution. Note that if the assumption of normality holds, the median calculated as suggested reverts to the mean. However, in the presence of outliers that skew the distribution of some variants, the median would be by definition more robust. We showed that the trade-off between HTS parameters, training set size and variant library distribution is a complex process that can potentially be both hardware- and problem-specific. In some situations, increasing the library size may be advantageous, while in others, optimizing the HTS parameters could be more beneficial. Rather than providing a one-size-fits-all solution, we aimed to equip researchers working with discretized HTS data with additional theoretical and practical tools. These tools are intended to help scientists better understand how binning their fitness values through FACS interval censoring could potentially impact experimental results, and thereby affect ML performance of models trained on such data.

## Acknowledgments and Disclosure of Funding

This work was supported by the University of Basel, the Swiss Federal Institute of Technology in Zürich (ETH Zürich) and the Swiss Nanoscience Institute (SNI, project P1802). The authors express their gratitude to Paolo Minelli for his insightful comments. The authors declare that some text was edited using ChatGPT (https://chat.openai.com). The editing focused solely on rearranging, enhancing, or correcting the syntax of existing sentences without creating new content. Portions of this work were performed at sciCORE scientific computing center at the University of Basel (https://scicore.unibas.ch).

