## Supporting Information for "Impact of Interval Censoring on Data Accuracy and Machine Learning Performance in Biological High-Throughput Screening"

### A Code and Data Availability Statement

The data and code produced for this work, including the documentation associated, are available free of charge on DOI:10.5281/zenodo.13840800 and github.com/Nash-Lab/HMLC.

### B Assumptions

#### B.1 Evenly Distributed and Constant Gates

We assumed that the gates were evenly distributed and with constant gate widths.

#### B.2 Normal Distribution

We assumed that the continuous fitness distribution of each variant was normally distributed.

#### B.3 Constant Standard Deviation

We assumed that each variant had the same continuous standard deviation ($\sigma_{y,i} = \sigma_y$).

#### B.4 Censored Standard Error

We assumed the validity of the central limit theorem (CLT) to estimate the standard error of each censored distribution ($\sigma_{\bar{y},i} = \sigma_y / \sqrt{N_{s,i}}$).

#### B.5 Combinatorial Variant Library Distribution

The combinatorial library variant population was assumed to follow a perfect binomial distribution.

#### B.6 Validation Set

While we changed the approximation levels (censor and/or sampling) of the training data, we used only the continuous means ($\mu_y$) to validate (hyperparameter optimization) and to test the models employed (LCs).

*
†

### C Materials and Methods

#### C.1 Theoretical distribution of a single variant ($N_s = \infty$)

We generated the censored mean ($\mu_z$, Fig. 2B) and censored standard deviation ($\sigma_z$, Fig. 2C) data of a theoretical distribution of a single gaussian changing the normalized gate width ($h/\sigma_y$) and normalized gate position ($z_0/h$) using Eq. S1 and Eq. S2.

$$\mu_z = \sum_{z \in \mathcal{H}} p_{gate}\left(z + h/2\right) \cdot \left(z + h/2\right) \tag{S1}$$

$$\sigma_z = \sum_{z \in \mathcal{H}} p_{gate}\left(z + h/2\right) \cdot \left(z + h/2 - \mu_z\right)^2 \tag{S2}$$

Where $\mu_y$ and $\sigma_y$ corresponded respectively to the mean and standard deviation of the continuous data distribution, and $\mathcal{H}$ was the ensemble of all gate edge coordinates.

The probability of each bin ($p_{gate}$) was calculated using Eq. S3.

$$p_{gate}(z + h/2) = \Phi\left(\frac{z + h - \mu_y}{\sigma_y}\right) - \Phi\left(\frac{z - \mu_y}{\sigma_y}\right) \tag{S3}$$

With $\Phi\left(\cdot\right)$ being the cumulative distribution function of the normal distribution (Eq. S4), which is closely related to the error function ($erf$).

$$\Phi\left(\frac{z - \mu_y}{\sigma_y}\right) = \frac{1}{2}\left[1 + erf\left(\frac{z - \mu_y}{\sigma_y\sqrt{2}}\right)\right] \tag{S4}$$

Moreover, we calculated the Shannon entropy ($H$, Fig. S1B) using Eq. S5 and Kullback–Leibler Divergence ($D_{KL}$, Fig. S1A) Eq. S6.

$$H(p_{gate}) = -\sum_{bin} p_{gate} \cdot \log_e\left(p_{gate}\right) \tag{S5}$$

$$D_{KL}\left(p_y || p_{z^*}\right) = \sum_{z \in \mathcal{H}} \int_{z^*=z}^{z+h} \int_{y=z}^{z+h} p_y \cdot \log_e\left(\frac{p_y}{p_{z^*}}\right) dydz^* = h(p_{z^*}) - h(p_y) \tag{S6}$$

Where $h$ was the differential entropy (Eq. S7) and $p_{z^*}$ was the continuous ("histogram"-based) version of the discrete distribution of $p_z$.

$$h\left(p_y\right) = -\int_y p_y \cdot \log_e\left(p_y\right) dy \tag{S7}$$

We utilized $p_{z^*}$ instead of $p_z$ to ensure that the Kullback–Leibler Divergence converged to zero in the limit of an infinitesimal small gate width (Eq. S8).

$$\lim_{h \to 0} D_{KL} = 0 \tag{S8}$$

#### C.2 Realistic distribution of a single variant ($N_s < \infty$)

During the calculation of $Bias$ and $Variance$ we marginalized the $z_0$ variable by calculating its expected value (Eq. S9).

$$\mathbb{E}_{z_0}\left[f\left(z_0\right)\right] = \int_{z_0=-0.5}^{0.5} z_0 \cdot f\left(z_0\right) dz_0 \tag{S9}$$

Due to symmetries, absolute values, and to avoid any problems in the numerical integration, Eq. S9 was further simplified to Eq. S10.

$$\mathbb{E}_{z_0}\left[f\left(z_0\right)\right] = 2\int_{z_0=0}^{0.5} z_0 \cdot f\left(z_0\right) dz_0 \tag{S10}$$

The equidistant points used in Fig. 3, 4, S3, S4, S5, S6 and S7 (red and blue lines) were calculated using the arc length formula (Eq. S11).

$$s = \int_a^b \sqrt{1 + \left(\frac{dy}{dx}\right)^2} dx \tag{S11}$$

Where $s$ was the arc length, and $a$ and $b$ were the integration limits. Moreover, the derivative of $y$ in respect of $x$ ($dy/dx$) was estimated using two cubic splines.

#### C.3 Realistic distribution of multiple variants

To obtain a library of noise-free labelled synthetic datapoints for this study, we relied on the ones generated in our previous work [1]. This was based on calculating the binding energies between the mutagenized library generated from a subset of the human fibrinogen beta chain (*FFFSARG*, $n = 7$) and the SdrG protein from S. epidermidis [4] using EvoDesign physical Energy Function (EvoEF) [3, 2]. To reduce the complexity of the input space and to ensure the full coverage of all possible mutations, we restricted the vocabulary of possible mutations to the ones already present in the WT sequence ($v = 5$). This included Alanine (A), Phenylalanine (F), Glycine (G), Arginine (R) and Serine (S). The total number of combinations per mutation (individuals and cumulative) were summarized in Tab. S1.

The individual frequency of each mutant in the in-silico libraries used in this work was calculated accordingly to a binomial distribution (Eq. 8). We explored two scenarios. In the first one, we further simplified the binomial distribution to the uniform distribution (Eq. 9) by setting the probability of the wild-type amino acids to one over the vocabulary size ($p_{WT} = 1/5$). In the second one, we set $p_{WT}$ by maximizing the information of the variants with a number of mutations equal or smaller than three (Eq. 10 and Eq. S12)

$$p_{WT}^{\#} := \underset{p_{WT}\in[0,1]}{\arg\max} \sum_i H\left(\frac{P_{bin}\left(x_i | m \in (0,\, 3]\,,\, n=7,\, v=5\right)}{\sum_i P_{bin}\left(x_i | m \in (0,\, 3]\,,\, n=7,\, v=5\right)}\right) \approx 0.69 \tag{S12}$$

After this, we calculated the probability of each variant with less than four mutations ($p\left(x_i | m \leq 3\right)$) following Eq. 8. Additionally, we assumed that the screened library was limited to up to triple mutants and we further normalized $p\left(x_i | m \leq 3\right)$ accordingly to Eq. S13.

$$p_{sl}\left(x_i | m \leq 3\right) = \frac{p\left(x_i | m \leq 3\right)}{\sum_i p\left(x_i | m \leq 3\right)} \tag{S13}$$

Where $p_{sl}\left(x_i | m \leq 3\right)$ was the normalized probability for each variant in the screened library (the ones presenting less than four mutations).

We then calculated the number of samples drawn for each variant ($N_{s,i}$) based on the desired average number of sample ($\bar{N}_s$) using Eq. S14.

$$N_{s,i} = p_{sl}\left(x_i | m \leq 3\right) \cdot \bar{N}_s \cdot \sum_{m=0}^{3} \binom{n}{m} \cdot (v-1)^m \Big|_{n=7,\, v=5} \tag{S14}$$

Note that here, in contrast to a real world scenario, $N_{s,i}$ was not restricted to integer values. This fact did not limit our sampling approach, based on CLT, but instead allowed us to cover all variants in the library, which would not be possible if we included a rounding step on $N_{s,i}$. The continuous averages for each variant ($\bar{y}_i$) were then sampled using Eq. S15.

$$\bar{y}_i \sim \mathcal{N}\left(\mu_{y,i} = f_{EvoEF}\left(x_i\right) \,,\, \sigma_{y,i} = \frac{\sigma_y}{N_{s,i}}\right) \tag{S15}$$

Where $\mu_{y,i}$ was the continuous mean of variant $x_i$, calculated using EvoEF function, and $\sigma_y$ was the general continuous standard deviation and it was set to half of the standard deviation calculated using all continuous means of the screened library (Eq. S16).

$$\sigma_y = \frac{1}{2}\sigma\left(\mu_{y,i}\left(x_i | m \leq 3\right)\right)\Big|_{n=7,\, v=5} \approx 1.7 \tag{S16}$$

Finally, the censored averages for each variant ($\bar{z}_i$) were sampled utilizing Eq. S17.

$$\bar{z}_i \sim \mathcal{N}\left(\mu_{z,i} \,,\, \frac{\sigma_{z,i}}{N_{s,i}}\right) \tag{S17}$$

Where $\mu_{z,i}$ and $\sigma_{z,i}$ were calculated respectively with Eq. S1 and Eq. S2. The gates were positioned to include all screened variants accordingly to the set described in Eq. S18.

$$\mathcal{H} = \left\{y \in \mathbb{R} | \, y = min\left(\mu_{y,i}^{i|m\leq3}\right) - 8\sigma_y + \theta \cdot h,\, \theta \in \mathbb{N}_0,\, y < max\left(\mu_{y,i}^{i|m\leq3}\right) + 8\sigma_y\right\} \tag{S18}$$

Where $h$ was the desired gate width.

#### C.4 ML performance on reconstructing the continuous distribution from censored training data

To test the ML performances, we used the methods already established in our previous work [1].

We encoded all sequences using one-hot encoding (OHE) at an amino-acid level. This choice was justified by the simplicity of such encoding and its broad use in biochemical studies with the aim of predicting proteins properties.

As ML models we utilized kernel ridge regression (KRR) [5] for its easy interpretability and training process, for its wide use in predictive chemistry, especially in the context of predicting various quantum properties, and for its closed form solution (Eq.S19).

$$\alpha_i = \sum_{j\,|\,m\leq3} \left[\left(K\left(x_i, x_j\right) + \lambda I\right)^{-1}\right]_{i\,|\,m\leq3,\, j} \mu_{y,j} \tag{S19}$$

Where $\alpha_i$ enclosed all trained parameters, $K\left(\cdot\right)$ was the Laplacian shift-invariant kernel (Eq. S20), $\lambda$ was a regularization hyperparameter, and $x_{i\,|\,m\leq3}$ corresponded to all sequences in the screened library (with less than four mutations), equivalent to the training set. The matrix inversion was computed using Cholesky decomposition.

$$K_{ij} = K\left(x_i, x_j\right) = exp\left(-\frac{||x_i - x_j||_1}{\sigma}\right) \tag{S20}$$

Where $\sigma$ was the second hyperparameter.

We optimized both hyperparameters via grid search by calculating the mean absolute error (MAE) on a validation set (see below), using two logarithmic scales with equally spaced points.

The validation and test sets comprised 8960 and 21504 data points, respectively, exclusively drawn from variants containing four or more mutations. As previously discussed, the size-variable training set was limited to variants with up to three mutations, with sorting based on the number of mutations present, referred as mutant-based shuffling. This setup mirrors a typical wet-lab experimental scenario, where a library is generated through methods such as error-prone Polymerase Chain Reaction (ep-PCR) or similar techniques, resulting in libraries that follow the binomial distribution previously described. In this context, the ML task was designed to extrapolate the ideal fitness values ($\mu_{y,i}$) of highly mutated variants (occupying a larger sequence space) based on censored data derived from low-order mutants ($\bar{z}_j$). This approach closely simulates the experimental challenge of predicting the behavior of more complex variants using information from simpler, less mutated ones.

Finally, the mutant-based normalized LCs were reported using a combinatorial scale (Eq. S21).

$$N^{norm} = \underset{\hat{m}\in[0,n]\subset\mathbb{R}}{\arg\min} \left( N - \sum_{p\in S(\hat{m})} \frac{\Gamma(n+1)\cdot(v-1)^p}{\Gamma(p+1)\cdot\Gamma(n-p+1)} \right) \tag{S21}$$

Where $N_{norm}$ and $N$ were respectively the normalized and non-normalized training set sizes, $n$ was the length of the peptide ($n = 7$), $v$ was the vocabulary size ($v = 5$), $\Gamma(\cdot)$ was the gamma function, and $p$ was defined over the set $S$ (Eq. S22)

$$S = \{p \in \mathbb{R} \,|\, p = \hat{m} - i,\ i \in \mathbb{N}_0,\ i \leq \hat{m} + 1\} \tag{S22}$$

Note that, due to the mutant-based shuffling of the training set, the normalized training set size ($N^{train}_{norm}$) was closely connected with the number of mutants enclosed within its datapoints. For examples, $N^{train}_{norm}$ equal to zero corresponded to only the WT, $N^{train}_{norm}$ equal to one corresponded to the WT and all single mutants and so on.

To generate a single LC for each combination of HTS parameters screened here, we trained 100 different models combining 10 different mutant-based shuffling and 10 noise sampling (Eq. S15 and Eq. S17) and we calculated the median of those 100 learning curves.

### D Supporting Figures

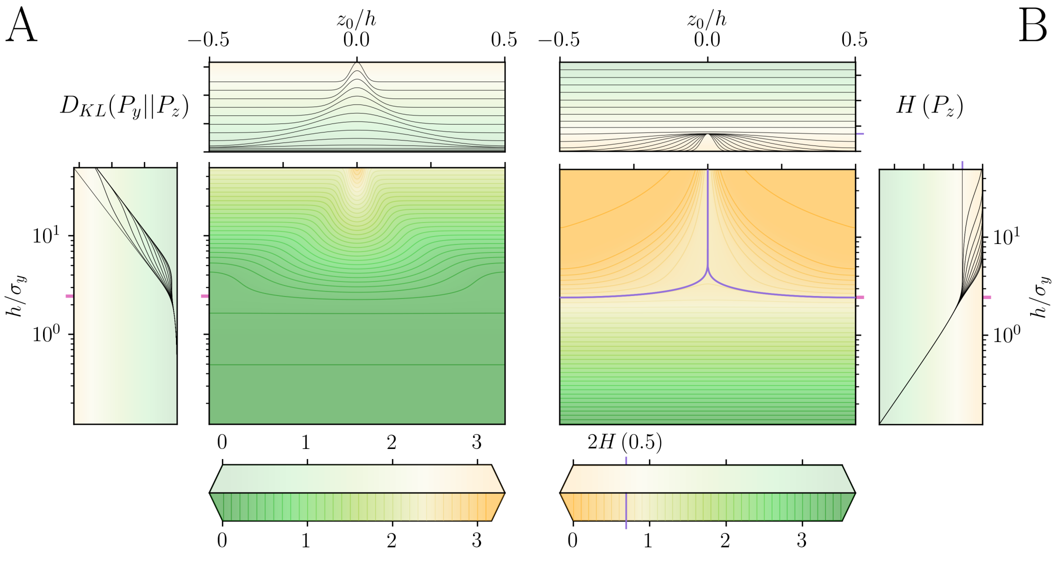


Figure S1: Impact of interval censoring on entropy-related measures, changing normalized gate width ($h/\sigma_y$) and normalized position ($z_0/h$). (A) Kullback–Leibler Divergence ($D_{KL}$) between continuous ($P_y$) and censored ($P_z$) distributions. (B) Shannon Entropy. The peak at high gate widths is reported as a purple line and corresponds to the entropy of a uniform binary distribution.

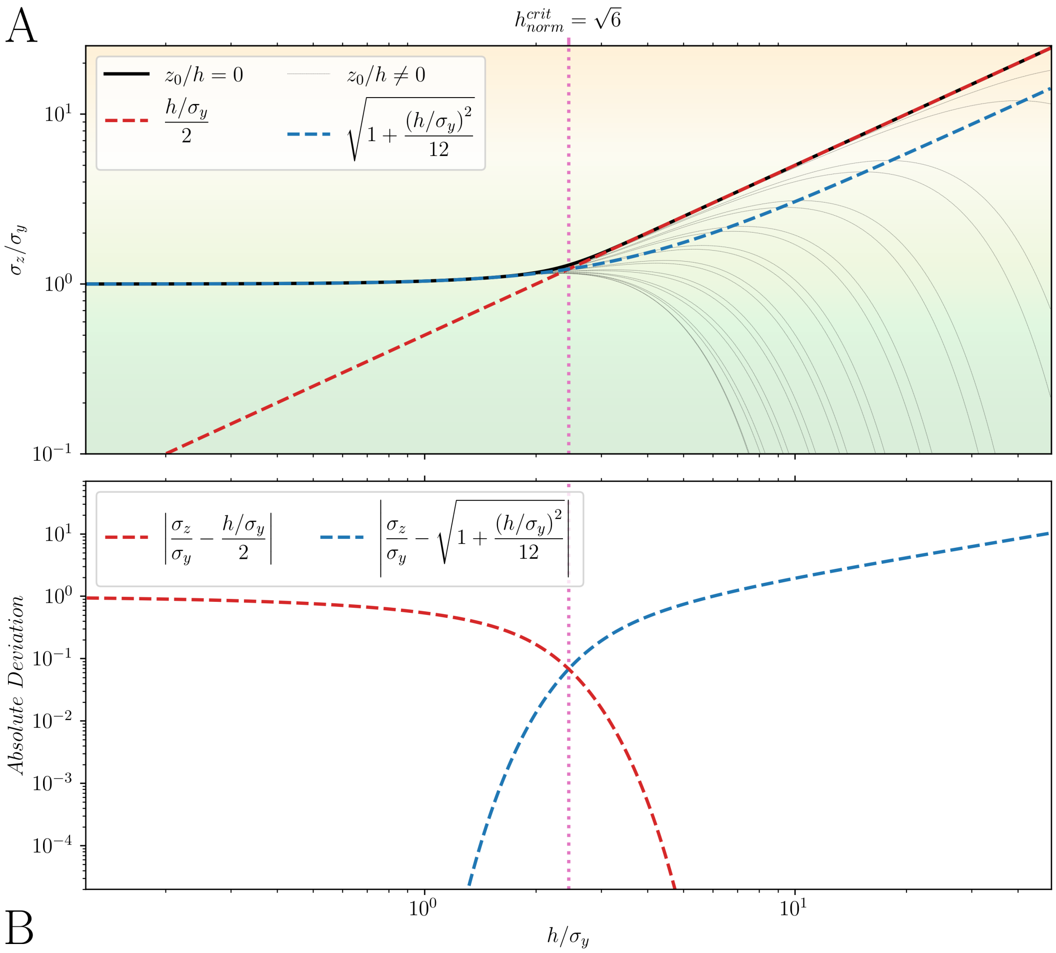


Figure S2: Derivation of normalized critical gates width ($h_{norm}^{crit}$). (A) Normalized censored standard deviation ($\sigma_z/\sigma_y$) versus normalized gates width ($h/\sigma_y$). The case where the continuous population was centered between two gates ($z_0 = 0$) is reported as a bold solid line. Other cases ($z_0 \neq 0$) are reported as thin dotted black lines. (B) Absolute difference between $\sigma_z/\sigma_y$ and the approximations (showed as red and blue lines). The normalized critical gate width was reported as a magenta dotted line.

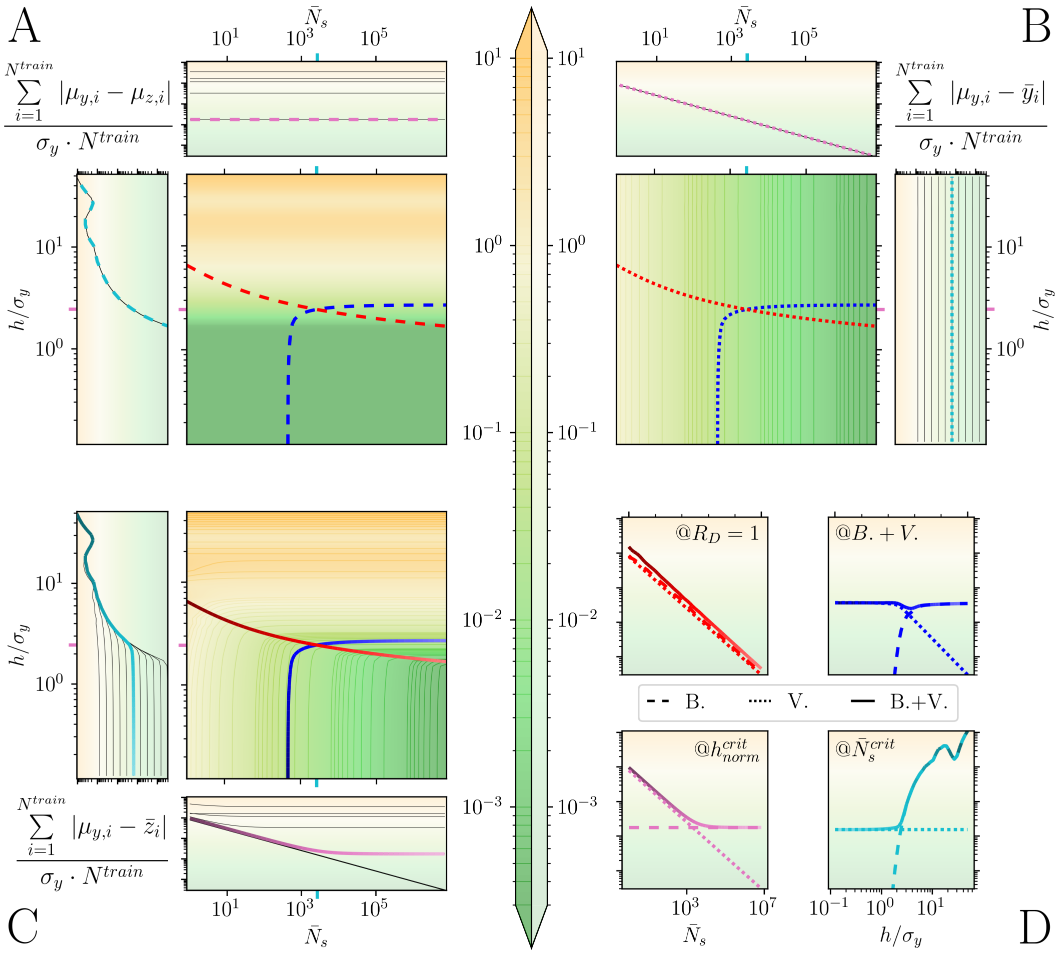


Figure S3: Impact of approximation on multiple variants (pooled library) at different fitness levels, changing the normalized gate width ($h/\sigma_y$) and the average number of samples ($\bar{N}_s$). Normalized MAEs between continuous means ($\mu_{y,i}$) and (A) censored means ($\mu_{z,i}$); (B) continuous averages ($\bar{y}_i$); and (C) censored averages ($\bar{z}_i$). (D) Contour cross-sections at different critical parameters: $R_D = 1$ (red), constant $B. + V.$ (blue), $h^{crit}_{norm}$ (magenta), and $N^{crit}_s$ (cyan). The colormap is shared between all panels.

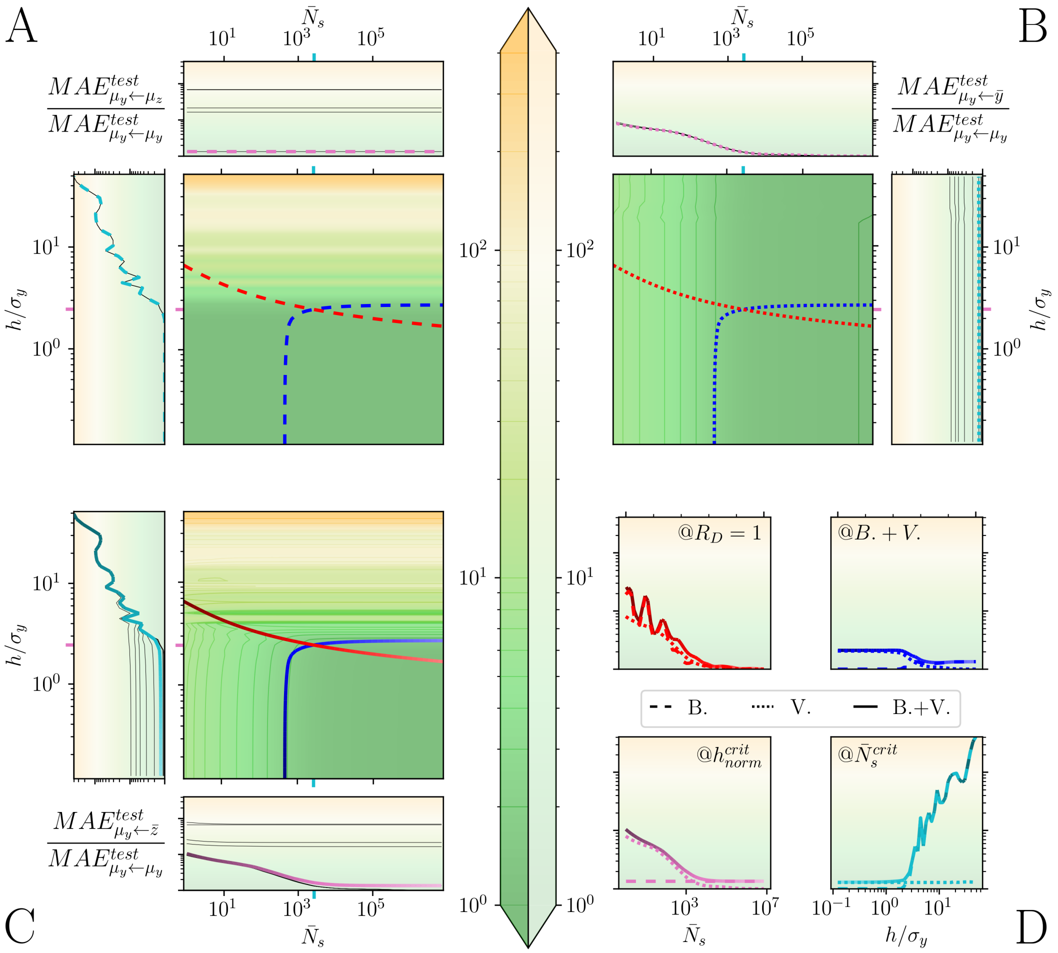


Figure S4: Impact of interval censoring on ML models trained with WT, single, double and triple mutants ($m = N^{train}_{norm} = 3$) changing the normalized gate width ($h/\sigma_y$) and the average number of samples ($\bar{N}_s$). Normalized MAEs of models trained with: (A) censored means ($\mu_z$); (B) continuous averages ($\bar{y}$); and (C) censored averages ($\bar{z}$). (D) Contour cross-sections at different critical parameters: $R_D = 1$ (red), constant $B. + V.$ (blue), $h^{crit}_{norm}$ (magenta), and $N^{crit}_s$ (cyan). The color scale shown in the middle applies uniformly across all panels.

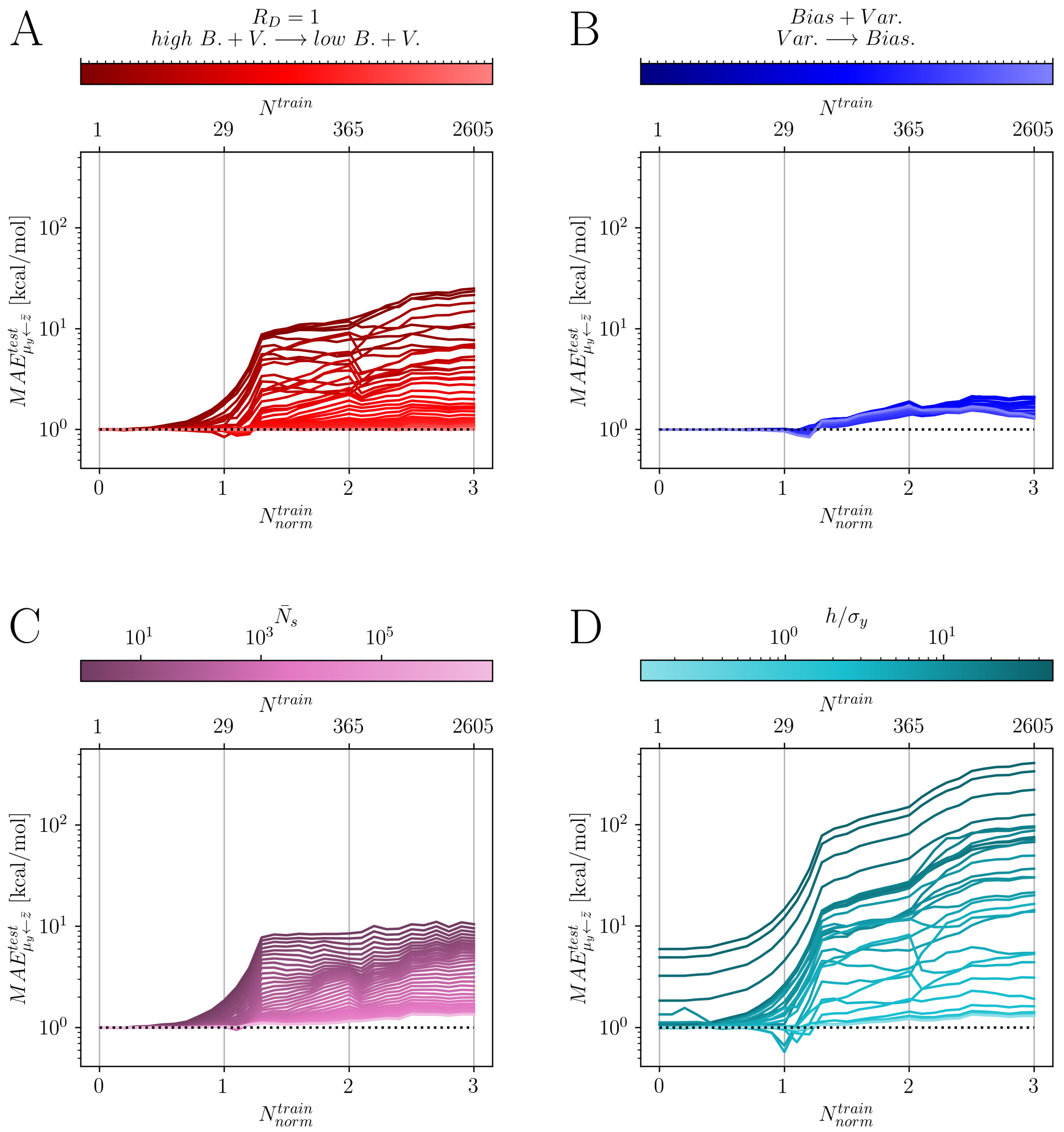


Figure S5: Impact of interval censoring on LCs ratio of models trained with up to triple mutants ($N^{train}_{norm} \in [0-3]$) as a function of the mutant-based normalized number of training points ($N^{train}_{norm}$), and using a uniform distributed combinatorial library ($P_{WT} = 0.2$). The plots illustrate the test performance under different conditions: (A) $R_D = 1$ with a gradient from high $B+V$ to low $B+V$ (red), (B) constant $B+V$ with varying $B$-$V$ contributions (blue), (C) constant normalized gate width ($h/\sigma_y = h^{crit}_{norm}$) with varying average sample size ($\bar{N}_s$, magenta), (D) constant average sample size ($\bar{N}_s = \bar{N}^{crit}_s$) with varying normalized gate width ($h/\sigma_y$, cyan).

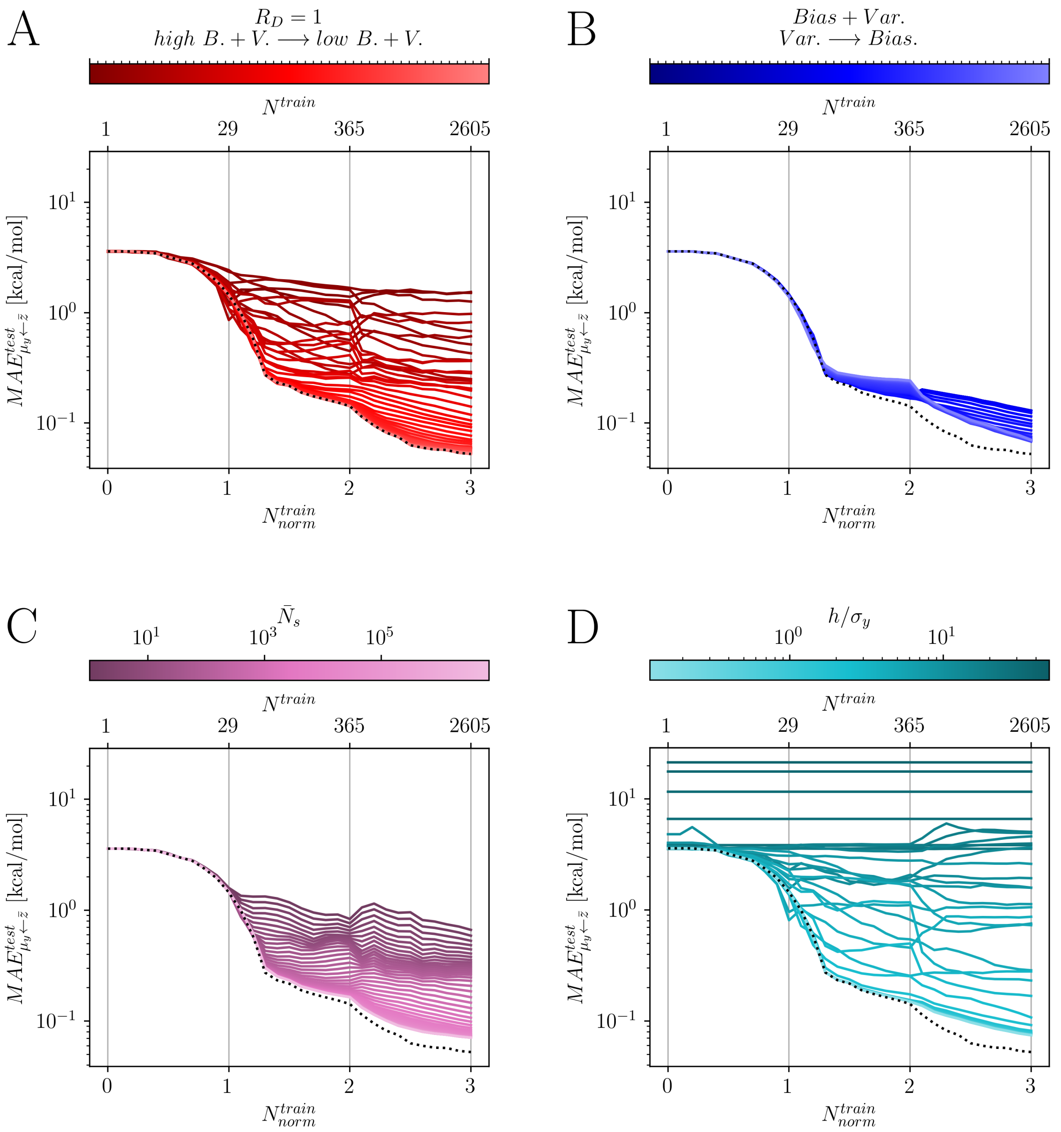


Figure S6: Impact of interval censoring on LCs of models trained with up to triple mutants ($N^{train}_{norm} \in [0-3]$) as a function of the mutant-based normalized number of training points ($N^{train}_{norm}$), and using a binomial distributed combinatorial library ($P_{WT} \approx 0.69$). The plots illustrate the test performance under different conditions: (A) $R_D = 1$ with a gradient from high $B+V$ to low $B+V$ (red), (B) constant $B+V$ with varying $B$-$V$ contributions (blue), (C) constant normalized gate width ($h/\sigma_y = h^{crit}_{norm}$) with varying average sample size ($\bar{N}_s$, magenta), (D) constant average sample size ($\bar{N}_s = \bar{N}^{crit}_s$) with varying normalized gate width ($h/\sigma_y$, cyan).

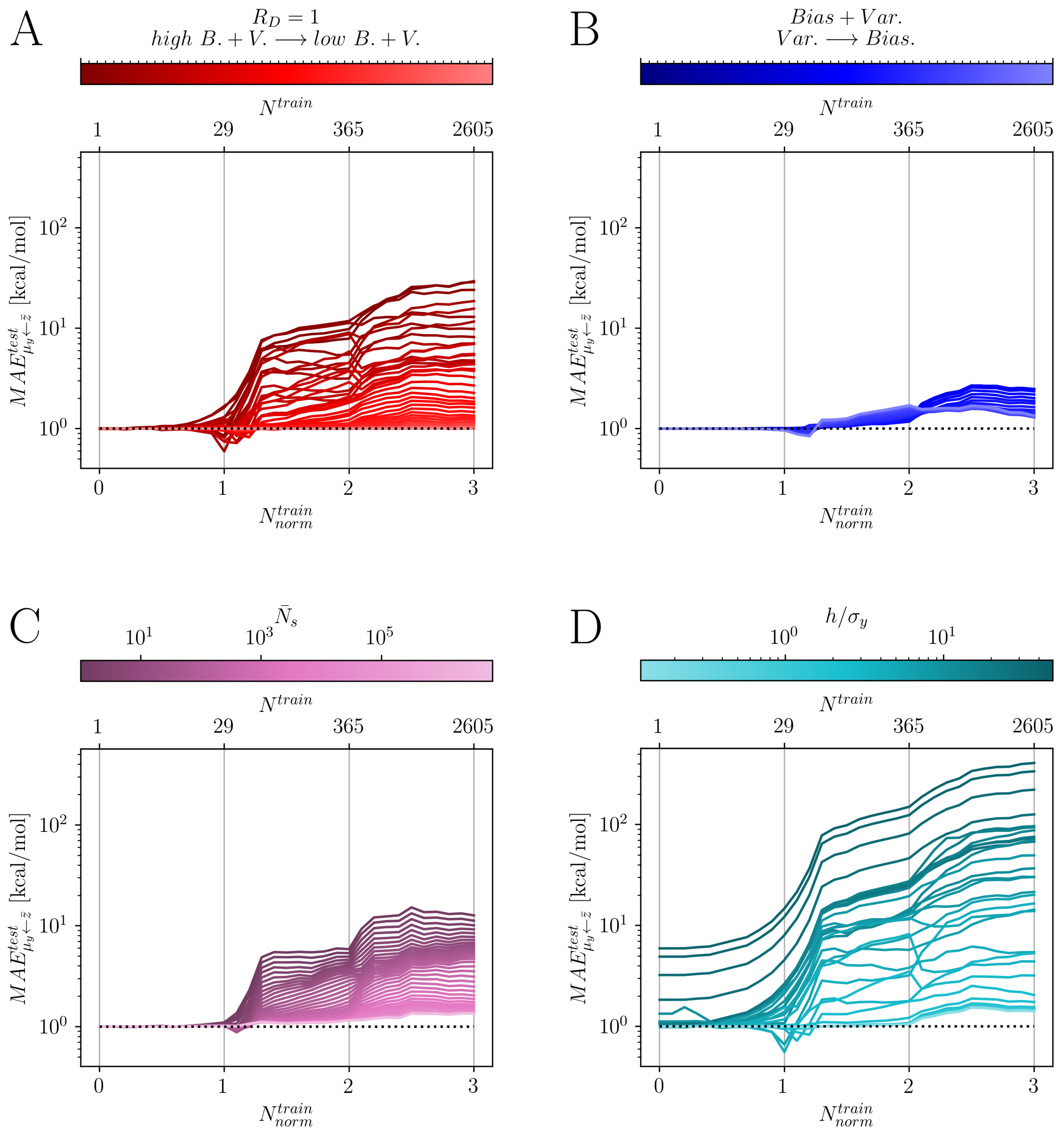


Figure S7: Impact of interval censoring on LCs ratio of models trained with up to triple mutants ($N^{train}_{norm} \in [0-3]$) as a function of the mutant-based normalized number of training points ($N^{train}_{norm}$), and using a binomial distributed combinatorial library ($P_{WT} \approx 0.69$). The plots illustrate the test performance under different conditions: (A) $R_D = 1$ with a gradient from high $B+V$ to low $B+V$ (red), (B) constant $B+V$ with varying $B$-$V$ contributions (blue), (C) constant normalized gate width ($h/\sigma_y = h^{crit}_{norm}$) with varying average sample size ($\bar{N}_s$, magenta), (D) constant average sample size ($\bar{N}_s = \bar{N}^{crit}_s$) with varying normalized gate width ($h/\sigma_y$, cyan).

### E Supporting Tables

Table S1: Size of mutagenized databases generated. The top part refers to the specific mutations (WT, single, double, etc.), while the bottom corresponds to the cumulative number of mutations (WT, WT+single, WT+single+double, etc.).

| **Database** | $m$=0 | $m$=1 | $m$=2 | $m$=3 | $m$=4 | $m$=5 | $m$=6 | $m$=7 |
|---|---|---|---|---|---|---|---|---|
| Individual | 1 | 28 | 336 | 2240 | 8960 | 21504 | 28672 | 16384 |
| Cumulative | 1 | 29 | 365 | 2605 | 11565 | 33069 | 61741 | 78125 |

Table S2: HTS Parameters, with their values (or ranges) and explanation. Values separated by a colon (:) indicate a range spanning between those extremes. For some parameters (*) we reported approximated values.

| Parameter | Value(s) | Unit | Explanation |
|---|---|---|---|
| $h/\sigma_y$ | $\sqrt{6}/20 : \sqrt{6} \cdot 20$ | - | Normalized gate width. |
| $z_0/h$ | -0.5 : 0.5 | - | Normalized distance between the true fitness / continuous means ($\mu_{y,i}$) and the closest gate boundary. |
| $\bar{N}_s$ | 1 : 7067989* | - | Average number of samples. In an experimental scenario, this is analogous to the average number of cells sorted per variant $i$. |
| $p_{WT}$ | 0.2 & 0.69* | - | Probability of wild type amino acid in the mutagenized library at each position. |
| $WT$ | *FFFSARG* | - | Wild type sequence (WT). This was used as a starting point to build the mutagenized library. |
| $n$ | 7 | - | Length of the WT sequence / mutagenized library sequences. |
| $v$ | 5 | - | Vocabulary size of possible mutations to build the mutagenized library from WT. This included Alanine (A), Phenylalanine (F), Glycine (G), Arginine (R) and Serine (S). |
| $\mu_{y,i}$ | as $f(x_i)$ | kcal/mol | True fitness values. One for each entry of the mutagenized library ($x_i$). |
| $f(x_i)$ | -38.97 : -5.42 | kcal/mol | Fitness function. It estimates the binding energy of each entry of the mutagenized library ($x_i$) with a target protein (SdrG) using EvoEF [3, 2]. |
| $\sigma_y$ | 1.7* | kcal/mol | Standard deviation of continuous fitness. Equal for each entry of the mutagenized library ($x_i$). |

Table S3: Levels of approximation of fitness

| Parameter | Explanation |
|---|---|
| $\mu_y$ | Continuous mean(s) / true fitness value(s). Fitness value(s) in a scenario where infinite samples were drawn from a continuous distribution. |
| $\mu_z$ | Censored mean(s). Fitness value(s) in a scenario where infinite samples are drawn from a censored distribution (due to sorting in gates) of fitness. It depends only on the gate width ($h$). |
| $\bar{y}$ | Continuous average(s). Fitness value(s) in a scenario where finite samples were drawn from a continuous distribution. It depends only on the number of samples ($N_s$). |
| $\bar{z}$ | Censored average(s) / real fitness data from sorting experiment(s). Fitness value(s) in a scenario where finite samples were drawn from a censored distribution (due to sorting in gates). It depends on gate width ($h$) and the number of samples ($N_s$). |

[5] Carl Edward Rasmussen and Christopher K. I. Williams. *Gaussian Processes for Machine Learning*. The MIT Press, 2006.
